# Investigating bacteria-phage interaction dynamics using droplet-based technology

**DOI:** 10.1101/2023.07.14.549014

**Authors:** Nela Nikolic, Vasileios Anagnostidis, Anuj Tiwari, Remy Chait, Fabrice Gielen

**Affiliations:** Living Systems Institute, University of Exeter, Exeter, UK; Department of Physics and Astronomy, Faculty of Environment, Science and Economy, University of Exeter, Exeter, UK; Translational Research Exchange @ Exeter, University of Exeter, Exeter, UK; Department of Biosciences, Faculty of Health and Life Sciences, University of Exeter, Exeter, UK

**Keywords:** *Escherichia coli*, phage, bacterial growth, phage-induced lysis, phage therapy, droplet microfluidics, time-lapse microscopy

## Abstract

An alarming rise in antimicrobial resistance worldwide has spurred efforts into the search for alternatives to antibiotic treatments. The use of bacteriophages, bacterial viruses harmless to humans, represents a promising approach with potential to treat bacterial infections (phage therapy). Recent advances in microscopy-based single-cell techniques have allowed researchers to develop new quantitative approaches for assessing the interactions between bacteria and phages, especially the ability of phages to eradicate bacterial pathogen populations. Here we combine droplet microfluidics with fluorescence time-lapse microscopy to characterize the growth and lysis dynamics of the bacterium *Escherichia coli* confined in droplets when challenged with phage. We investigated phages that promote lysis of infected *E. coli* cells, specifically, a phage species with DNA genome, T7 (*Escherichia virus T7*) and two phage species with RNA genomes, MS2 (*Emesvirus zinderi*) and Qβ (*Qubevirus durum*). Our microfluidic trapping device generated and immobilized picoliter-sized droplets, enabling stable imaging of bacterial growth and lysis in a temperature-controlled setup. Temporal information on bacterial population size was recorded for up to 25 hours, allowing us to determine growth rates of bacterial populations helping us uncover the extent and speed of phage infection. In the long-term, the development of novel microfluidic and single-cell techniques will expedite research towards understanding the genetic and molecular basis of rapid phage-induced lysis, preempting bacterial resistance to phages and ultimately identifying key factors influencing the success of phage therapy.

## INTRODUCTION

Bacterial infections were the second leading cause of death globally prior the COVID pandemic [Ikuta et al 2022]. According to the World Health Organization, over 50% of life-threatening bacterial infections are caused by pathogens resistant to one or more antibiotics, with *Escherichia coli* being the leading antibiotic-resistant pathogen responsible for the most deaths in 2019 [WHO 2019, Ikuta et al 2022, Murray et al 2022]. In particular, *E. coli* pathogenic strains involved in urinary tract infection (UTI) and food poisoning are major public health concerns worldwide. Antibiotic treatments are typically prescribed against UTIs, however *E. coli* can be found in the patient urinary tract weeks after antibiotic treatment, increasing the chance of relapse and chronic infections [Klein & Hultgren 2020]. UTI is the source for more than 50% of bloodstream infection (bacteremia) cases, and over 40% of the UTI-causing strains are resistant to some of the antimicrobials commonly used [Allocati et al 2013, Bontenet al 2021, McCowan et al 2022, Murray et al 2022]. Antibiotic treatments against foodborne pathogens, Shiga toxin-producing *E. coli* strains are usually not recommended because the antibiotic-induced bacterial SOS response increases the Shiga toxin production and release, which damages host cells and can lead to severe disease outcomes [Mühlen & Dersch 2020].

One option to consider when designing treatments against infections caused by multi-drug resistant or toxin-producing bacterial pathogens, would be to employ natural killers of bacteria, their viruses – phages, to be part of the antimicrobial treatment, referred to as phage therapy [Abedon 2019, Kortright et al 2019, Abedon et al 2021, Venturini et al 2022, Maimaiti et al 2022, Igler 2022, Strathdee et al 2023, Petrovic Fabijan et al 2023]. Phages are selective towards the bacterial host they infect, and usually narrowly target a single bacterial species or even a specific strain within the species. Phages are currently used as therapeutics for humans only in compassionate cases, however, phage therapy is already in veterinary use in livestock and other animals [McCallin et al 2019]. Even though phages can interact with host immune systems, they do not seem to trigger a strong immune response, and are thus far considered nontoxic to humans and animals [Nale & Clokie 2021, Champagne-Jorgensen et al 2023]. Phage monotherapy and phage cocktail (a mixture of two or more phage species) are therefore promising alternatives for treating infections caused by clinically relevant bacterial pathogens where antibiotics are of no benefit or should even be completely avoided [Dedrick et al 2019, Yang et al 2020]. Phages can also be administered in conjunction with conventional antibiotics, with several cases known where phage-antibiotic combinations eradicated infections more efficiently than the antibiotic alone [Van Nieuwenhuyse et al 2022, Suh et al 2022]. Employing phage therapy as a novel clinical practice would require access to appropriate phage collections [Gibson et al 2019, Maffei et al 2021], and methodologies to rapidly determine the efficacy of phage-based treatments to eradicate pathogen populations and the likelihood of developing phage resistance.

Recent metagenomic studies have revealed novel phage sequences and substantially expanded their number in public databases, which subsequently led to restructuring of phage taxonomy [Lefkowitz et al 2017, Walker et al 2022]. This shed light on phage genomic diversity, however without addressing their host range. Current bioinformatics methods fall short of predicting host range of phages solely from their genomics data and therefore cannot be used on their own [Zrelovs et al 2020]. Instead, rapid and quantitative functional assays are required to evaluate phage infectivity and lytic activity against candidate bacterial hosts, especially clinical bacterial isolates. The standard plaque assays for phage enumeration and detection often involve demanding multistep protocols and long culturing times, and provide only limited throughput. On the other hand, high-throughput phage characterization can be performed but requires expensive and complex robotic systems [Chory et al 2021].

Techniques able to obtain data at the single-cell level such as high-resolution microscopy will enable rapid screening of many microenvironments in parallel. In addition, time-lapse microscopy data can help gather extensive temporal information on bacteria-phage dynamics. This would therefore accelerate phage characterization procedures by quickly quantifying bacterial growth reduction and bacteriolysis at the single-cell level [Attrill et al 2021, Nikolic et al 2023].

Current developments in combining microscopy with droplet microfluidic technologies have paved the way towards high-throughput quantitative analysis of living systems. Droplet technology enables controlled encapsulation of cells into water-in-oil microreactors [Matuła et al 2020, Xu et al 2020, Sun et al 2023]. Every reactor, typically of picoliter volume, represents a unique environment in which cells can proliferate. Droplet-based technology has to date been successfully applied to multiple research fields, including the directed evolution of enzymes or single-cell -omics protocols [Gielen et al 2016, Gielen et al 2018, Anagnostidis et al 2020, Gantz et al 2023]. Coupling of microfluidic-based systems with high-resolution microscopy enables a wide diversity of phenotypic screens, for instance cellular secretions, antibody production or the study of cell population growth [Liu et al 2016, Kleine-Brüggeney et al 2019, Rutkauskaite et al 2022].

Several microfluidic-based microscopy setups have been successfully employed for monitoring fluorescently-labeled bacteria in droplets at the single-cell and population levels, using fluorescence signal as a proxy for number of bacterial cells or population size [Leung et al 2012, Huang et al 2015, Kaushik et al 2017, Mahler et al 2018, Postek et al 2018, Barizien et al 2019, Pratt et al 2019, Hsu et al 2019, Taylor et al 2022]. These studies have shown that microfluidic droplet technology can be used not only to quantify the growth of various bacterial species and microbial communities in different conditions, but also to test bacterial susceptibility to a range of antimicrobial compounds. In addition, droplet technology has been applied to develop phage detection and enumeration tools [Tjhung et al 2014, Yu et al 2014]. However, these methods have not been adapted to study bacteria-phage interactions, and in particular have not explored whether lysis by phages can be successfully quantified from single-plane fluorescence maps. In our study, we developed a novel microfluidic device that generates anchored droplets populated with *E. coli* and their phage, and embed air cavities used for precise autofocus. This collectively allowed for long-term evaluation of bacterial population dynamics across multiple droplets, enabling quantitative screening of phages as therapeutics to treat bacterial infections.

## MATERIALS AND METHODS

### Microfluidic device

Our microfluidic chips were fabricated using two-layer soft lithography processes (**Figure 1A**) [Bentley et al 2022]. Even though the chips contained one, two or three rows of traps located between rows of pillars, all traps had the same size and geometry (**Supplementary Figure S1**). The pillars contained square cavities filled with air during chip operation and used for the autofocus function. The height of the bottom flow channel was 7.5 μm. The top layer had circular traps of diameter of 60 μm and was 7.5 μm high, such that the overall height of the droplets was 15 μm. Uncured polydimethylsiloxane (PDMS) consisting of a 10:1 polymer to cross-linker mixture (Sylgard 184) was poured onto the master, degassed, and baked at 70°C for at least 4 hours. Before this curing time at 70°C, a piece of 0.15 mm thick glass cover slip was immersed into the liquid PDMS and manually aligned above the trapping array. This method minimized droplet evaporation into the PDMS (**Supplementary Figure S2**). Following master mold fabrication, PDMS chips were plasma bonded (Diener Zepto) on thin cover slip glass (thickness 0.15 mm) to enable high-resolution imaging. Hydrophobic surface treatment was performed after bonding by flushing with 1% (v/v) trichloro(1H,1H,2H,2H-perfluorooctyl)silane (Merck) in HFE-7500 and, subsequently, placed in a 65°C oven for 30 min.

**Figure 1.**
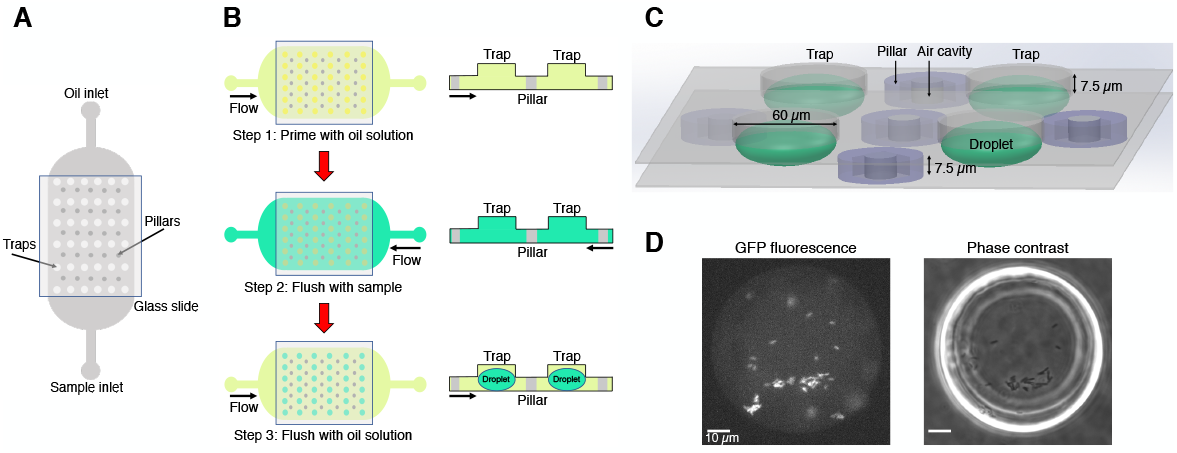
Microfluidics formation of anchored droplets for the study of bacteria-phage interactions. **A)** The device schematic shows the top view of the microfluidic chip with two inlets, used for flushing oil and samples through the whole device respectively. The trapping array was covered with a thin glass cover slip (see **Methods** for details). **B)** The three-step self-digitization method was utilized to generate droplets on-chip. **C)** Three-dimensional schematic depicts droplets formed within a chip, along with pillars embedding air cavities, and the size of traps. **D)** The fluorescence and phase contrast images show droplets that contain *gfp*-carrying *E. coli*. Scale bars represent 10 μm.

### Droplet generation

The microfluidic chip was placed on an epifluorescence microscope equipped with an automated stage. A self-digitization water-in-oil emulsion method was followed to make droplets (**Figure 1B**). First, the entire microfluidic chip was manually primed with a fluorinated oil (HFE-7500, Fluorochem Ltd) solution supplemented with 1% w/w fluorosurfactant (008, RAN Biotechnologies) using a syringe. The main reason for using a surfactant in the oil solution is to reduce the interfacial tension between the sample and the oil phase. This facilitates the digitization process and prevents droplet coalescence [Cohen et al 2010]. The sample containing either bacteria or bacteria-phage mixture was then flushed through the chip using the opposite inlet to replace the HFE oil solution. The oil solution was subsequently flushed again through the chip forming droplets of volume 28 picoliters (assuming elliptical spheres).

### Fluorescence time-lapse microscopy

Time-lapse imaging data were acquired using a temperature-incubated, μManager-controlled Olympus IX83 fluorescence microscope equipped with an LED light source (Lumencor SpectraX) and an automated stage (Marzhauser) [Edelstein et al 2014, Chait et al 2017]. The incubation box (cellVivo) temperature was maintained at 30°C for all assays. Fluorescence and phase contrast images were acquired with an sCMOS camera (Hamamatsu Orca Flash4v3) every 3 min through a 40× objective (UPLFLN40XPH, Evident Olympus). Up to six positions were recorded in each time-lapse, capturing four to six droplets per field of view. Air cavities in pillars of the microfluidic chip aided software-based autofocus (on 640 nm reflection images) performed prior to acquiring each image set (**Figure 1C**), as described in [Chait et al 2017]. We then recorded a phase contrast image to confirm that traps are populated with sample-containing droplets, and a GFP fluorescence image (to quantify fluorescence-based population size) at a focal plane mid-way through the droplets at each position (**Figure 1D**). We set exposure time to 20 ms for phase images, and 100 ms was the exposure time to image GFP (excitation 460-480 nm, emission 495-540 nm, dichroic mirror 490 nm).

### Image analysis

Raw fluorescence images were divided by a shading corrector image to account for inhomogeneity of GFP excitation illumination across the field of view, as described in [Chait et al 2017]. To obtain the shading corrector, we took a median projection through a stack of GFP images from distinct locations on a glass cover slip overlaid with 50% (w/v) fluorescein [Model et al 2001]. We then normalized this corrector image by dividing by its median pixel value. After shading correction, we performed background subtraction by using ‘mask’ images. Masks were made by finding the thresholds in each frame for which the background fluorescence value, i.e. signal not coming from fluorescent cells within droplets, corresponded to 0. Then, each shading-corrected image was multiplied with the respective mask image for each recorded frame. This final image was used to set a circular region of interest (ROI) around each droplet. We extracted mean fluorescence intensity per ROI corresponding to mean droplet fluorescence intensity. For the analysis, we excluded frames in which the device was out of focus. All image analysis routines were done in the open-source image processing package Fiji (ImageJ) [Schindelin et al 2012], and more details can be found in **Supplementary Methods**.

### Phages, strains and cultivation

One phage species with DNA genome, T7 (*Escherichia virus T7*) and two phage species with RNA genomes, MS2 (*Emesvirus zinderi*) and Qβ (*Qubevirus durum*) were used in this study, together with *E. coli* strains BW25113, W1485, and the derivatives of K-12 MG1655 harboring the constitutively expressed *gfp* reporter gene encoding for Superfolder GFP, with all details described in **Supplementary Table S1**. The F plasmid from strain W1485 was introduced by conjugation into strain TB193, now annotated as strain TB193 F+. Only strains that harbor the F plasmid can be infected by MS2 and Qβ, as RNA phage start infection by adsorbing to bacterial F-pili encoded on the F plasmid. Phage T7 can infect all listed *E. coli* strains. Bacterial cultures were grown in LB medium (1% tryptone, 0.5% yeast extract, 1% NaCl), which is a nutrient-rich medium. Frozen glycerol clones were streaked on LB agar plates to obtain single colonies. A single colony was used to inoculate overnight batch cultures shaking at 230 rpm. All overnight incubations and batch cultivations prior to phage infection were carried at 37°C. Bacterial cultures were supplemented with 0.01% glucose and 2 mM CaCl_2_ prior to adding RNA phage for infection experiments (addition of divalent ions facilitates phage adsorption [Rappaport 1965, Ács et al 2020]). LB medium was supplemented with 10 μg/ml chloramphenicol for selection after conjugation.

### Sample preparation for droplet assays

Overnight cultures of an *E. coli* strain were diluted 1 to 100 into 4 ml of fresh LB medium for 3 h at 37°C, to obtain exponentially growing cultures at ∼5*10^8^ bacterial cells/ml. Bacterial cultures were then supplemented with 0.01% glucose and 2 mM CaCl_2_. For phage infection experiments, bacterial cultures were mixed with phage at a specific multiplicity of infection (MOI, the ratio of phage particles to bacterial cells in a culture). Samples were loaded into the microfluidic device, and the exact time from preparing the sample to starting the time-lapse microscopy experiment was noted (typically, 11-14 min). An aliquot of the bacterial culture was taken just before starting the microscopy experiment (or in the case of infection experiments, just before adding phage) to determine the initial bacterial population size. Serial dilutions of the culture aliquot were plated on LB agar plates and incubated at 37°C for 24 h. Bacterial population size was determined by measuring the density of colony forming units CFU, as: CFU/ml= n_colonies_ / (dilution factor * V_diluted culture_).

### Phage lysate preparation

Overnight bacterial cultures of *E. coli* host propagation strain (BW25113 for phage T7, W1485 for phages MS2 and Qβ) were diluted 1 to 100 into LB medium. After 4 h, exponentially growing cultures were inoculated with a single plaque in 4 ml of phage soft agar (1% tryptone, 0.1% yeast extract, 0.01% glucose, 0.8% NaCl, 2 mM CaCl_2_, 0.7% agar; kept at 50°C), plated on phage plates (1% tryptone, 0.1% yeast extract, 0.01% glucose, 0.8% NaCl, 2 mM CaCl_2_, 1% agar), and incubated overnight at 37°C for 20 h [Nikolic et al 2023]. The following day, soft agar containing plaques was removed into a 50 ml-Falcon tube with 12 ml of SM buffer (0.1 M NaCl, 8 mM MgSO_4_·7H_2_O, 50 mM Tris-Cl pH 7.5, 0.01% gelatin). Falcon tubes were centrifuged for 15 min at 4000 g. The supernatant containing phage lysate was filtered twice through a 0.22 μm-filter into a new tube. Phage lysates were kept at 4°C, and their phage titer was determined by plaque spotting assays described below.

### Plaque spotting assays

Overnight cultures of the *E. coli* host strain were diluted 1 to 1000 into LB medium. After 5.5 h, 200 μl of exponentially growing bacterial cultures were added to 4 ml of phage soft agar, and plated on phage plates [Nikolic et al 2023]. After 2-3 min, serial dilutions of the phage lysate were spotted on top of the soft agar containing bacterial host, and the plates were incubated overnight for 20 h, at 37°C for phages MS2 and Qβ, or at room temperature for phage T7. Phage titer was determined by measuring the density of plaque-forming units PFU, as: PFU/ml= n_plaques_ / (dilution factor * V_diluted phage_).

### Plate-reader experiments

Overnight *E. coli* cultures were diluted 1 to 100 into LB medium. After 3 h of cultivation, the cultures were supplemented with 0.01% glucose and 2 mM CaCl_2_. Aliquots of the cultures were put into wells of a 96-well microplate, and each well was either infected with phage at the specific MOI, or the same volume of SM buffer was added to uninfected cultures. All samples were then diluted 1 to 10 into fresh LB medium, having final volume of 180 μl in each well. Four independent replicate cultures were analyzed in experiments with phage T7, and five independent replicates were analyzed in infection experiments with MS2 and Qβ. Growth of the cultures was recorded at 30°C, every 4 min for 800 min in total, as absorbance at 600 nm A_600_, with shaking prior to each measurement (CLARIOstar Plate Reader, BMG Labtech).

### Analysis of growth and lysis

For each growth curve, we employed the Microsoft Excel function *slope* from ln-transformed measurements over a sliding window of 40 min, i.e. over 11 time points in plate-reader assays or 14 time points in droplet assays. Bacterial growth rate μ was then defined as the maximum slope value for each growth curve, within the period *t*= 36-120 min for droplet experiments, and *t*= 20-120 min for plate-reader assays. In addition, growth rates μ of bacterial populations challenged with RNA phage were determined within the period *t*= 38-200 min for droplet experiments, and *t*= 24-200 min for plate-reader assays. Doubling times (plate-reader assays) or fluorescence-based doubling times (droplet assays) corresponded to ln2/μ. To mark the transition from initial bacterial growth to the onset of bacteriolysis, inflection points were evaluated for each growth curve from slopes calculated over a sliding window of 40 min, by identifying the time point where slope value changed from positive to negative. The middle time point of the sliding window was noted to be the inflection point. To evaluate differences between different conditions or different strains we used two-tailed two-sample heteroscedastic Student’s *t*-tests.

## RESULTS

### Adapting droplet technologies for phage assays

We optimized the layout of a microfluidic device for easy self-digitization, enabling formation of stable droplet-environments and steady microscopy imaging routine. The microfluidic chip utilized in our experiments had two inlets, one for the oil phase and the other one for the aqueous samples, with a central large trapping chamber embedding dozens of traps and pillars (**Figure 1A**). A glass cover slip was immersed in the uncured PDMS and cured in place on top of the trapping area to minimize droplet evaporation and ensure the stability of droplets during the course of time-lapse experiments (**Supplementary Figure S2**). The chip was placed on an automated fluorescence microscopy imaging platform in an enclosed environmental box set to 30°C. We employed a three-step process to generate droplets on-chip following the self-digitization emulsion method, which means that emulsions were formed by consecutive flowing of aqueous and oil phases into the same chip (**Figure 1B**). In our study, the aqueous phase was the sample of either a sole bacterial culture, or a mixture of phage and susceptible bacteria cultivated in growth medium. The chips were first primed with the solution of oil containing surfactant, and the sample was then flushed through the chip using the opposite inlet so that the sample could fill all traps. The oil solution was then flushed again and droplets were passively formed within the traps. Interfacial tension and density difference between aqueous samples and oil generated anchored droplets that had the diameter of the designed traps (**Supplementary Figure S1**). In our experiments, droplets were formed in 96% of traps within the field of view, while the remaining 4% were either empty traps or traps populated with more than one droplet (**Supplementary Dataset**). Moreover, each microfluidic chip consisted of rows of traps followed by rows of pillars (**Figure 1C**). Air cavities were included at the center of pillars to identify them and provide fiducials at the glass interface to aid software-based autofocusing performed using 640 nm illumination prior to imaging at each time point and location [Chait et al 2017] **(Supplementary Figure S1**). Stable autofocusing enabled steady image acquisition from several pre-selected positions on the chip per each time point, and ensured reproducible imaging despite stage motion (**Figure 1D**). The small height of the chamber (15 μm) ensured accurate quantification of overall fluorescence produced by the cells. Finally, we utilized a rapid and simple image analysis routine from a widely-used open-source image analysis software platform Fiji (ImageJ) to quantify mean fluorescence intensity of each droplet (see **Supplementary Methods**). Altogether, we developed a device and workflow for the on-chip droplet generation and stable multidroplet microscopy image acquisition dedicated to the long-term monitoring of bacterial growth and lysis in droplets.

### Evaluating the efficacy of phage-induced lysis in droplets

We first assessed how droplet-environments support the growth of *Escherichia coli* carrying chromosomally encoded, constitutively expressed *superfolder gfp*. An exponentially growing *E. coli* culture was loaded into a primed microfluidic device subsequently flushed with oil to form bacteria containing-droplets within traps. We monitored the increase in GFP fluorescence coming specifically from all cells during the time-lapse microscopy experiment, as the proxy for bacterial population growth in droplets. Despite known stochastic fluctuations in *gfp* expression in clonal bacterial populations, a broadly linear correlation between integrated droplet fluorescence and the number of fluorescent bacterial cells in droplets has been reported previously [Barizien et al 2019, Taylor et al 2022]. For this and all subsequent droplet experiments, we aimed at loading 10 to 40 bacterial cells in each droplet, with the number of bacterial cells across droplets following a Poisson distribution [Collins et al 2015, Taylor et al 2022]. We measured a mean 30-fold GFP fluorescence increase of the droplet populations over 400 minutes of the time-lapse experiment (**Figure 2A, Supplementary Movie S1**). Our analysis showed that the mean fluorescence-based doubling time of bacterial populations in droplets was 24 ± 8 minutes (mean ± standard deviation). The mean doubling time of the same strain growing in a plate-reader was 34 ± 2 minutes (**Figure 2B**). Overall, our results indicate that *E. coli* populations can successfully grow inside droplet-environments within designed microfluidic devices, with a growth pattern comparable to bulk batch cultures.

**Figure 2.**
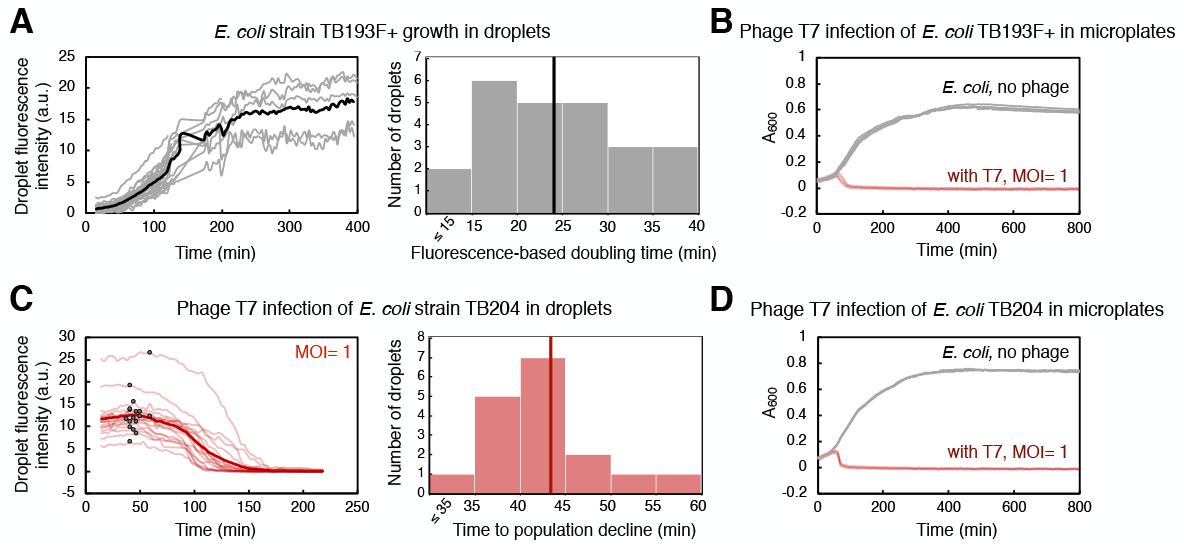
Bacterial growth and phage T7-induced bacterial population decline in droplets and in microplates. **A)** An exponentially growing culture of *E. coli* strain TB193 F+ harboring the *gfp* fluorescent reporter gene was loaded into a microfluidic device at 2.2*10^8^ CFU/ml (CFU, colony forming unit). GFP fluorescence intensity over time for 24 individual droplets is plotted in gray, and the black line is the mean value over all GFP fluorescence trajectories. The GFP signal of all droplets was collected for at least 120 min, and five individual droplets were followed until 400 min (see **Supplementary Methods** and **Supplementary Datasets**). The histogram shows the distribution of fluorescence-based doubling times of *E. coli* droplet populations growing in nutrient-rich medium, with the mean doubling time of 24 min (black line). **B)** Population size of TB193 F+ cultures, either infected with phage T7 or uninfected, was monitored in a plate-reader by recording absorbance at 600 nm (A_600_). Phage T7 was added to the cultures at time 0 and multiplicity of infection, MOI= 1 (4 independent replicate cultures, light red lines), or no phage was added (4 replicates, gray lines). **C)** *E. coli* strain TB204 harboring the *gfp* fluorescent reporter gene (6.3*10^8^ CFU/ml) was mixed with phage T7, at MOI= 1. Temporal information for GFP fluorescence intensity of 17 individual droplets is plotted in light red, and the thick red line is the mean value over all droplets. We determined the inflection point for each trajectory corresponding to the start of bacteriolysis in each droplet, i.e. the time point after which the bacterial droplet population size declines (indicated as black dots on light red lines). The histogram shows the distribution of times to population size decline, with the mean time of 43 min (thick red line). **D)** Population growth and lysis assays of *E. coli* strain TB204 were performed in a plate-reader, with 4 uninfected and 4 infected independent replicate cultures.

Next, we investigated whether we can utilize the same droplet-based setup to monitor the dynamics of bacterial populations challenged with phages. To this end, we mixed bacterial cultures of *gfp*-carrying *E. coli* with phage lysate prior to loading the sample into a microfluidic device. In all our experiments, we employed obligatory lytic phages, which are phages that promote bacteriolysis and kill infected bacterial cells. Subsequently, decline in bacterial population size due to lysis of bacterial cells, i.e. the biomass reduction within droplets, was detected as a decrease in overall GFP fluorescence. In the first set of experiments, we infected *E. coli* cultures with phage T7 at MOI= 1. T7 is a phage species with DNA genome, and is known for its short replication cycle (time from infection to the lysis of the host cell) of typically 15 minutes at 37°C, and 30 minutes at 30°C, and a high efficacy in killing *E. coli* cells [Jack et al 2019, Xu et al 2021]. By measuring GFP fluorescence, we detected a severe reduction in the *E. coli* population size during T7 infection in droplets. Our analysis indicated that on average 99.4% of an *E. coli* droplet population was killed 218 minutes after adding T7 phage (**Figure 2C**), with 0-2 cells per droplet remaining at the end of the time-lapse experiment (**Supplementary Movie S2, Supplementary Movie S3**). We determined the time point when infected bacteria in droplets had a population dynamics shifting from dominant growth to dominant lysis, by finding the inflection point of GFP fluorescence curves. The mean time to the population decline, i.e. to the start of significant bacteriolysis of droplet populations, was 43 ± 7 minutes. Growth assays in a plate-reader showed that the mean time to population decline during T7 infection was 46 ± 2 minutes, with the eradication of bacterial cultures, i.e. >99.9% of the population killed, happening after 165 ± 92 minutes (**Figure 2D**). In addition, time to T7-induced bacteriolysis was not significantly dependent on the presence of the *gfp* reporter gene in the *E. coli* genome (**Supplementary Figure S3**). The presence of the F plasmid slightly increased the time to lysis, by 19% on average, without significantly affecting the time to whole population eradication (**Supplementary Figure S3, Supplementary Table S2**). Overall, these experiments suggest that droplet-based technology can be utilized to analyze bacterial population growth dynamics in response to phage, and to quantify the efficacy of phages to kill bacterial populations.

### Investigating long-term interactions between bacteria and phages

We evaluated the ability of generally poorly characterized phages with RNA genomes to influence the growth of *E. coli* populations in droplets. MS2 (*Emesvirus zinderi*) and Qβ (*Qubevirus durum*) are the best known RNA phages, with longer phage replication cycle (typically 90 and 105 minutes at 37°C, respectively), different lysis machinery, and lesser efficacy in killing *E. coli* than DNA phage T7 [Rappaport 1965, Jenkins et al 1974, Woody and Cliver 1995, Bernhardt et al 2002, Tsukada et al 2009, Nikolic et al 2023]. *E. coli* were mixed with RNA phage lysates at MOI ∼10, to ensure a large excess of infective phage particles; specifically, MS2 lysate at MOI= 7, and Qβ lysate at MOI= 14. Droplets populated with bacteria-phage samples were then generated and trapped within a microfluidic device. Phage MS2 impeded bacterial growth in droplets, with fluorescence-based doubling time of droplet populations increasing more than 3-fold compared to uninfected populations, to 81 ± 16 minutes (**Figure 3A, Supplementary Movie S4**). Analysis of inflection points from GFP fluorescence curves indicated that droplet population size started to decline 374 ± 72 minutes after *E. coli* being challenged with MS2, in 63% of analyzed droplets. The remaining droplet populations either exhibited a reduced growth rate without population size decline, or resumed growth shortly after the initial population size decline (**Supplementary Dataset**). Overall, after 601 minutes of exposure to phage MS2, we measured a mean 2.8-fold GFP fluorescence increase of the droplet populations compared to the GFP fluorescence intensity at the beginning of the experiment. In addition, we measured the growth dynamics of the same *E. coli* strain challenged with phage MS2 at MOI= 10 in a plate-reader (**Figure 3B**). The population doubling time in a plate-reader was 46 ± 4 minutes during MS2 exposure, and the size of all plate-reader populations started to decrease 310 ± 45 minutes after adding phage MS2. Furthermore, we detected a reduction in bacterial growth after challenging *E. coli* with phage Qβ, with the fluorescence-based doubling time of 204 ± 110 minutes for infected populations in droplets (**Figure 3C, Supplementary Movie S5**). The start of decline of the droplet population size was detected 233 ± 18 minutes after adding phage Qβ, in 94% of analyzed droplets. At the end of the time-lapse experiment involving 1481 minutes of exposure to phage Qβ, we measured a mean 1.6-fold GFP fluorescence-decrease of the droplet populations. Similarly to the experiments with phage MS2, we measured growth dynamics of *E. coli* populations challenged with phage Qβ at MOI= 10 in a plate-reader (**Figure 3D**). The population doubling time during Qβ exposure was 43 ± 3 minutes, and decline in the size of all plate-reader populations was noticed 445 ± 17 minutes after *E. coli* being challenged with phage Qβ. Altogether, these results indicate that utilizing droplet-based technology can provide quantitative temporal information on how phages modulate bacterial growth rate and alter population size of their bacterial host over longer periods of time. Comparable phage-modulated bacterial growth dynamics between plate-reader and droplet experiments paves the way towards development of high-throughput multidroplet screening methods that can make bacteria-phage interaction analyses faster than standard microbiology methods.

**Figure 3.**
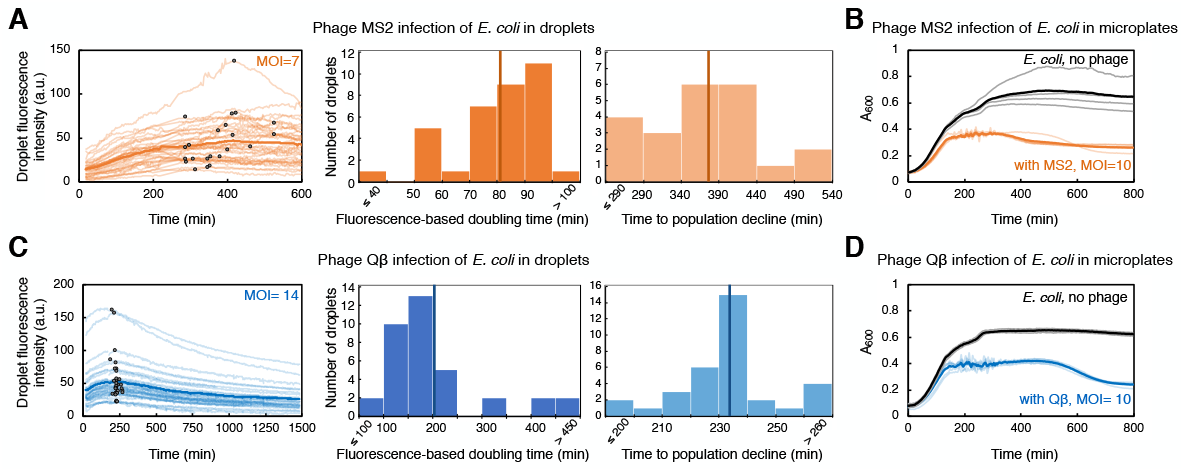
*E. coli* challenged with RNA phage in droplets and in microplates. **A)** *E. coli* strain TB193 F+ (5.9*10^8^ CFU/ml) was challenged with phage MS2, at MOI= 7. GFP fluorescence intensity of 35 individual droplets during MS2 exposure is plotted in bright orange (with inflection points depicted with black dots), and the dark orange line is the mean value across all droplets. The left histogram shows the distribution of fluorescence-based doubling times, which were estimated from the growth rate calculated during the first 200 min of the experiment. The mean value of 81 min is depicted with a dark orange line. The right histogram shows the distribution of times to the droplet population decline upon MS2 phage infection, inferred from inflection points of GFP fluorescence curves of 22 droplets, with the mean value of 374 min. **B)** The same *E. coli* strain was mixed with phage MS2 at MOI= 10, and bacterial growth and lysis were monitored in a plate-reader. Light orange depicts lysis curves of 5 independent replicates, with the mean value across all replicates depicted in dark orange. Growth curves of 4 uninfected replicate cultures are depicted in gray, with the mean value across all replicates depicted in black. **C)** *E. coli* culture (7.2*10^8^ CFU/ml) was challenged with phage Qβ, at MOI= 14. GFP fluorescence intensity recorded for 36 individual droplets during Qβ exposure is plotted in light blue (with inflection points indicated as black dots), and blue line is the mean value across all droplets. The left histogram shows the distribution of fluorescence-based doubling times, with the mean value of 204 min depicted with dark blue line. The right histogram shows the distribution of times to the droplet-population decline upon Qβ infection, inferred from inflection points of GFP fluorescence curves of 34 droplets, with the mean value of 233 min. **D)** The growth dynamics of *E. coli* challenged with phage Qβ at MOI= 10 was recorded in a plate-reader. Light blue presents lysis curves of 5 infected independent replicate cultures, with the mean value across all replicates shown in blue. Growth curves of 3 uninfected replicate cultures are depicted in gray, with the mean value depicted in black.

## DISCUSSION

We have established a novel methodology that couples droplet-based assays in a microfluidic device to time-lapse fluorescence microscopy for studying bacteria-phage interactions and population dynamics. The described droplet generation technique eliminates the need for complex and expensive equipment (e.g. pumps), making it an accessible and cost-effective method for generating and monitoring droplets on-chip. Furthermore, our methodology delivers a unique approach to detect changes in bacterial population size upon phage infection, in a temperature-controlled environment, ensuring the stability of droplets and steady image acquisition over long time periods. For the image analysis pipeline, we employed a set of basic tools freely available from an open-source platform, without implementing any additional custom-made scripts or closed-source software packages. We successfully quantified the speed (time to bacteriolysis) and extent of invasion (the fraction of lysed bacterial population and the bacterial growth reduction) from three different phage species attacking a common bacterial species (**Figure 4**). As such, this methodology has a great potential to allow us to routinely assess both phages that can be effective pathogen-killers and phages that can modulate the structure of host-associated commensal bacterial populations.

**Figure 4.**
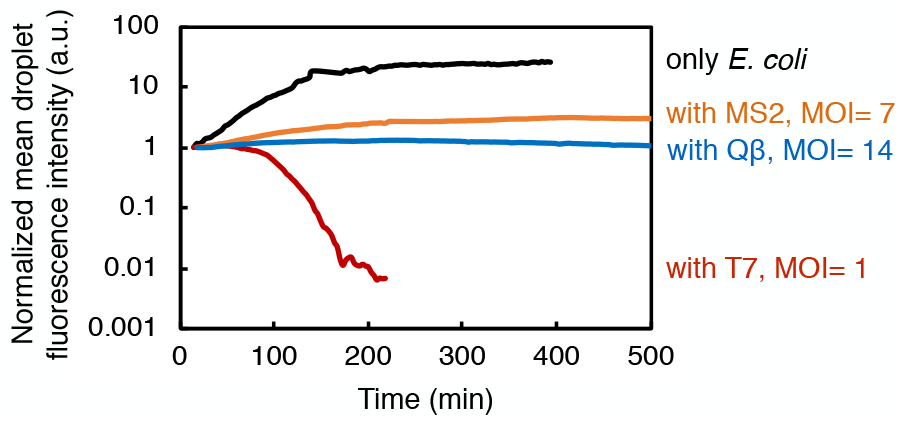
Summary of the results showing bacterial growth and lysis in droplets. Here presented are normalized mean values of droplet fluorescence intensities from all experiments. Overall, our study indicates the potential of droplet-based technology to quantify the bacterial population dynamics in response to phage exposure.

Our high-resolution imaging modality enabled quantification of fluorescence signals down to single-cell resolution provided cell numbers were low and cells were in, or close to, optimum focus. Even though some cells were out-of-focus, they still contributed to the final mean fluorescence intensity readout provided their GFP signal was above a threshold value (see **Methods** and **Supplementary Methods**). This issue was also mitigated by cell motility, which ensured consistent number of cells found in and out of focus across the time series. In addition, a stable autofocus function ensured data acquisition at the reproducible plane of focus for comparable datasets over the experiment duration. On the whole, absolute quantification of cell number was prevented mainly by autoaggregation (formation of bacterial clumps [Trunk et al 2018]) and the presence of cells far from the focal plane.

The established droplet-based methodology could be utilized to understand ecological and (co)evolutionary aspects of dynamics between bacteria and their phages, especially bacteria-phage dynamics at low bacterial population densities [van Houte et al 2016, Koskella et al 2022]. For instance, stable long-term time-lapse imaging of droplet-environments at a specific temperature could provide the optimal platform for investigating interactions between temperate phages and their bacterial hosts. Temperate phages, in addition to their lytic cycle, can also display a lysogenic life cycle by integrating their genome into the genome of their host. Droplet assays could help apprehend dynamics of switching from lysogenic to lytic cycle and *vice versa*, and provide insights into how this switch changes structure and behavior of the bacterial host population. In particular, the peptide signal-mediated short-range communication between phages, which controls lysogeny-lysis decision based on the bacterial host densities [Erez et al 2017, Aframian et al 2022], could be monitored in droplet-environments.

In addition to capturing quorum sensing molecules and secreted proteins, droplet-environments can allow bacterial cells to maintain native planktonic and aggregate lifestyles and form biofilms [Chang et al 2015]. The aggregate formation can be induced by various stressors [Trunk et al 2018, Cai 2020], and can occur even as a response to phage infection [Cai et al 2022]. Recent time-lapse microscopy studies of bacteria-phage dynamics employed microfluidic devices with confined habitats (so called mother machines), which support one-dimensional growth of bacterial cells preventing them from forming aggregates or biofilms [Attrill et al 2021, Nikolic et al 2023]. Our experiments showed that aggregates can be formed from single bacterial cells within a droplet (**Supplementary Movie S1**), thus enabling us to evaluate phage efficacy against individual cells as well as aggregates within the population.

Extension of the method to screening a larger number of droplets will provide more insights into stochastic processes at play when bacterial populations start growing from just few cells [Barizien et al 2019]. This could be done by designing devices with more traps in the field of view. With more droplet assays per time-lapse experiment, rare events corresponding to emergence of bacterial resistance (genetic basis) and tolerance (non-genetic basis) to phages could also be identified, and resilient bacteria extracted from the chip to determine the underlying mechanisms that shape bacterial antiphage defense strategies. In addition to bacterial strains that have chromosomally encoded or plasmid-based fluorescent gene reporters, our setup could be employed to analyze bacterial cultures of any strain stained with fluorescent dyes prior to phage infection [Yoon et al 2021]. Moreover, future work could utilize fluorescent dyes to label phages outside host cells [Egido et al 2023], to apprehend dynamics of both bacteria and phage numbers over time, and better understand the progression of phage infection. Finally, alternative single-cell methodologies that do not require bacteria to be fluorescently-labeled, would be useful to detect growth and phage-induced lysis of individual bacterial cells from brightfield microscopy images using deep learning-based algorithms [Howell et al 2022, Tiwari et al 2023]. To conclude, the development of novel population-level and single-cell tools and approaches will expedite research towards understanding what makes phages effective bacteria-killers and how their bacterial hosts can eventually mitigate phage invasion, highlighting the best phage candidates for phage therapy against specific bacterial diseases.

## DATA AVAILABILITY

All datasets generated in this study are available within the Supplementary Data (‘SupplementaryDatasets.zip’). Design files for microfluidic devices are deposited on DropBase (http://openwetware.org/wiki/DropBase).

## AUTHOR CONTRIBUTIONS

NN and FG designed the study; NN, VA, AT, and RC performed the experiments; NN did image and data analysis; NN, RC, and FG interpreted the data; NN and FG wrote the manuscript with input from all co-authors.

## ACKNOWLEDGEMENTS

The authors thank Lucy Witherall and Wolfram Möbius for sharing the phage T7 lysate and strain BW25113, and for their technical support, and Tobias Bergmiller for sharing bacterial strains. We acknowledge the LSI Technical Services Team at the University of Exeter and use of the Exeter Microfluidics Facility and Savchenko Centre for Nanoscience.

