## Supplementary Information for "Investigating bacteria-phage interaction dynamics using droplet-based technology"

**Supplementary Figures**
Figures S1-S3

**Supplementary Tables**
Tables S1-S2

**Supplementary Movies**
Movies S1-S5

**Supplementary Datasets**
Source data for figures and analysis

**Supplementary Methods**

**Supplementary References**

**SUPPLEMENTARY FIGURES**

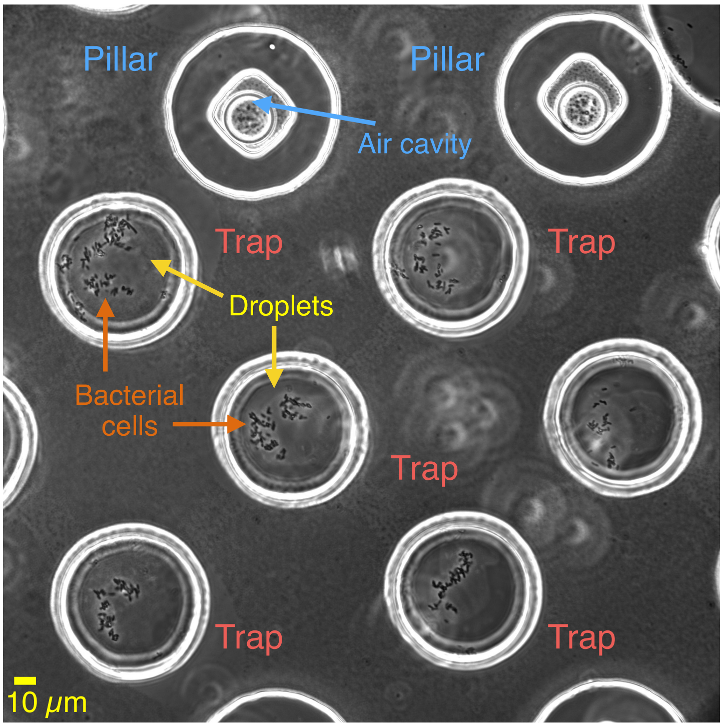


**Supplementary Figure S1.** Single position on the microfluidic chip.
The phase contrast microscopy image shows a field of view at 40x, with three rows of traps populated with bacteria-containing droplets, and with one top row of pillars. Each pillar has a cavity in the center, filled with air bubbles, to aid software-based autofocusing prior to image acquisition at each location and time point. Scale bar represents 10 μm.

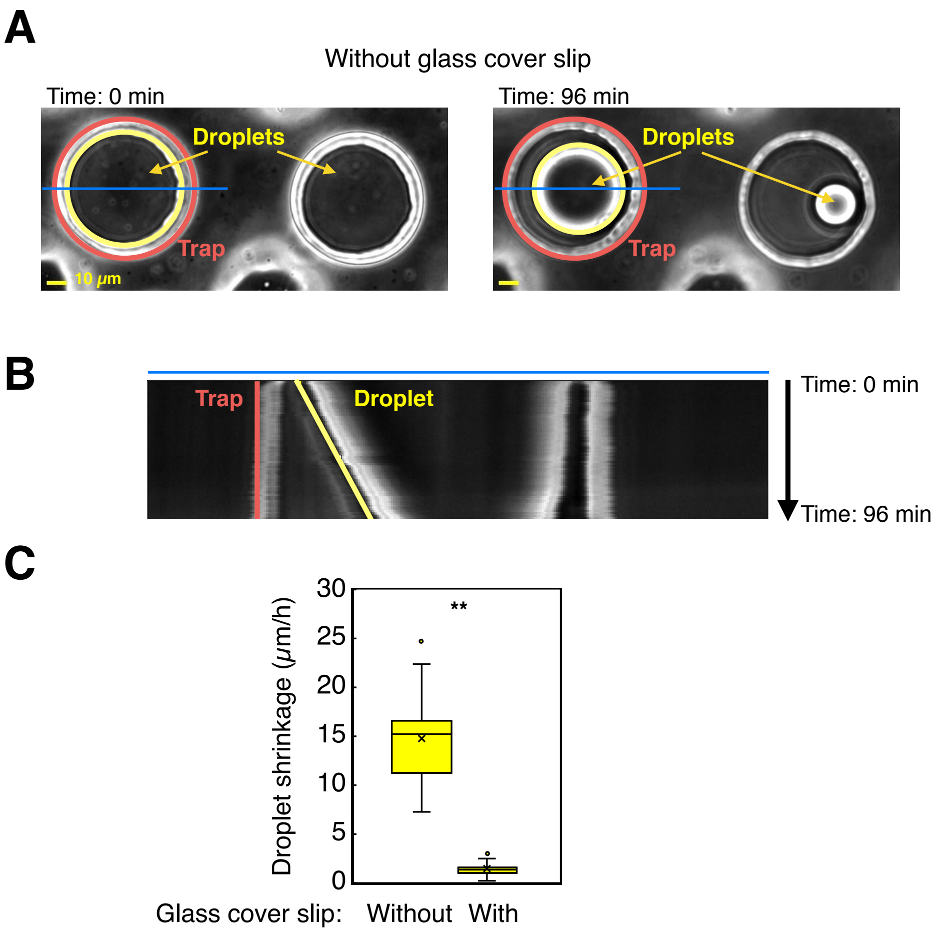


**Supplementary Figure S2.** Analysis of droplet shrinkage.

**A)** Phase contrast images from the beginning (*t*= 0 min) and the end (*t*= 96 min) of a time-lapse experiment show a substantial decrease in the droplet size when a glass cover slip was not placed on top of the trapping area of the microfluidic chip. Yellow circle depicts the droplet circumference, while the red circle depicts the trap wall. A blue line was drawn across the trap (to include the trap diameter), traversing the trap wall and droplet edge. Scale bar represents 10 µm. **B)** Kymographs were generated from time-lapse movies in Fiji (ImageJ), and used to plot the pixel intensity along a chosen line (i.e. blue line shown in **A)**) as a function of time. The phase contrast images were acquired every 1 min through a 20× objective (Olympus UPLFLN20X), over a period of 96 min. **C)** In the experiment without the glass cover slip on the top of the microfluidic chip, we analyzed 12 droplets, and in the experiment with the glass cover slip we analyzed 11 droplets. The diameter of each droplet was noted at *t*= 0 min and *t*= 96 min, and droplet shrinkage was calculated as the difference in the droplet diameter in the period of 96 min. Putting the glass cover slip on the chip substantially decreased the evaporation and thereby prevented the volume reduction of a droplet (*t*-test, *P*= 2e-06).

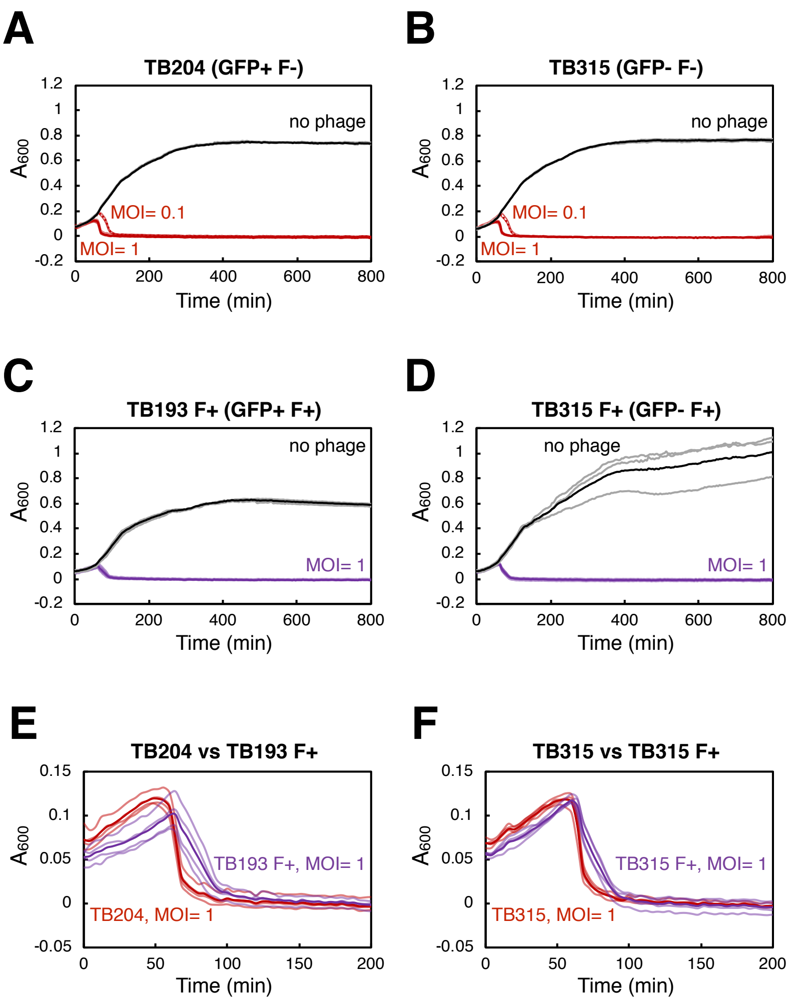


**Supplementary Figure S3.** T7 infection of *E. coli* strains in microplates.

An aliquot of exponentially growing cultures of *E. coli* strains **A)** TB204 (dataset partially represented in **Figure 2D**), **B)** TB315, **C)** TB193 F+ (dataset presented in **Figure 2B**), and **D)** TB315 F+, was mixed with phage T7 at MOI= 1 and MOI= 0.1, while the remaining cultures stayed uninfected. Bacterial population growth and lysis of 4 independent replicate cultures (3 uninfected independent replicates in panel **D)**) was monitored in a plate-reader every 4 min for 800 min in total. Light red or light purple depicts curves of infected replicate cultures, with the mean value across all replicates depicted in thick red or thick purple. Uninfected controls are presented in gray, with the mean value across all replicates depicted in black. In some panels, the growth and lysis differences between replicate cultures are so small that the replicate curves substantially overlap. Start of the population size decline (time to population lysis) was not dependent on the presence of the *gfp* reporter gene: TB204 vs TB315 *P*= 0.18 (*t*-test), TB193 F+ vs TB315 F+ *P*= 0.05. **E) and F)** Direct comparisons of the lysis curves from panels **A)** and **C)**, as well as from panels **B)** and **D)** respectively (zoomed in to the first 200 min of the experiment), indicate that time to population lysis was dependent on the presence of the F plasmid: TB204 vs TB193 F+ *P*= 0.0003, and TB315 vs TB315 F+ *P*= 0.014. However, time to the bacterial population clearance (eradication point, i.e. >99.9% of the population killed) was not dependent on the presence of the F plasmid: TB204 vs TB193 F+ *P*= 0.72, TB315 vs TB315 F+ *P*= 0.57.

### SUPPLEMENTARY TABLES

**Supplementary Table S1.** Phages and bacterial strains used in this study.

| Name | Relevant characteristics | Source or reference |
|---|---|---|
| *Escherichia* phage T7 | *Escherichia virus T7*<br>dsDNA bacteriophage | Wolfram Möbius Collection |
| *Enterobacteria* phage MS2 | *Emesvirus zinderi*<br>(+)ssRNA bacteriophage | DSM 13767, Leibniz Institute DSMZ-German Collection of Microorganisms and Cell Cultures |
| *Enterobacteria* phage Qβ | *Qubevirus durum*<br>(+)ssRNA bacteriophage | DSM 13768, Leibniz Institute DSMZ-German Collection of Microorganisms and Cell Cultures |
| MG1655 | *E. coli* K-12 wild-type<br>F-, λ-, *ilvG-, rfb-50, rph-1* | [Blattner et al 1997]<br>Tobias Bergmiller Strain Collection |
| W1485 | *E. coli* Lederberg W1485; F+ *met str*, T1s T6s λ-<br>Classical F+ donor in conjugation<br>Host strain for propagation of phages MS2 and Qβ | DSM 5695, Leibniz Institute DSMZ-German Collection of Microorganisms and Cell Cultures |
| BW25113 | *E. coli* K-12 strain<br>F-, λ-, Δ(*araD-araB)567*, Δ*lacZ4787*(::*rrnB-3*), *rph-1*, Δ(*rhaD-rhaB)568*, *hsdR514*<br>Host strain for propagation of phage T7 | [Datsenko & Wanner 2000]<br>Wolfram Möbius Collection |
| TB315 | MG1655 Δ*araA*::*kanR* | Tobias Bergmiller Strain Collection |
| TB315 F+ | F plasmid from W1485 introduced by conjugation in TB315 | [Nikolic et al 2023] |
| TB193 | MG1655 *att*P21::λ$P_R$-*gfp*::frt-*cat*<br>constitutive *gfp* expression<br>*gfp* encodes Superfolder GFP | Tobias Bergmiller Strain Collection |
| TB193 F+ | F plasmid from W1485 introduced by conjugation in TB193 | This study |
| TB204 | MG1655 *att*P21::λ$P_R$-*gfp*::frt | [Bergmiller et al 2017] |

**Supplementary Table S2.** Growth and T7-induced lysis of *E. coli* cultures in microplates.

| *E. coli* strain | Strain description | [a] Start of the population size decline (min), MOI= 1 | [b] Bacterial population clearance (min), MOI= 1 | Start of the population size decline (min), MOI= 0.1 | Bacterial population clearance (min), MOI= 0.1 | Doubling time of uninfected cultures (min) |
|---|---|---|---|---|---|---|
| TB204 | F- GFP+ | [*]46 ± 2 | 165 ± 92 | 65 ± 2 | 264 ± 81 | 36 ± 0 |
| TB315 | F- GFP- | 48 ± 0 | 139 ± 36 | 64 ± 0 | 208 ± 82 | 33 ± 1 |
| TB193 F+ | F+ GFP+ | 58 ± 2 | 186 ± 61 | not measured | not measured | 34 ± 2 |
| TB315 F+ | F+ GFP- | 54 ± 2 | 166 ± 80 | not measured | not measured | 34 ± 2 |

[a] inflection point
[b] eradication point
[*] mean ± standard deviation

### SUPPLEMENTARY MOVIES

**Movie S1.** Time-lapse of a population of *E. coli* growing in a droplet. (Scale bar corresponds to 10 µm.)
**Movie S2.** Time-lapse of *E. coli* cells in a droplet during exposure to phage T7, showing lysis of the entire droplet-population.
**Movie S3.** Time-lapse of *E. coli* cells in a droplet during exposure to phage T7, with 2 cells left at the end of the experiment.
**Movie S4.** Time-lapse of *E. coli* cells in a droplet during exposure to phage MS2.
**Movie S5.** Time-lapse of *E. coli* cells in a droplet during exposure to phage Qβ.

### SUPPLEMENTARY METHODS

#### Image and data analysis routine

Here we describe step by step pipelines for shading correction, background subtraction, ROI (region of interest) generation and image analysis, and data analysis.

**A. Shading correction** (specific to the objective employed and fluorescence), according to [Chait et al 2017]

Preparation of the shading corrector image:

1. Get 12 GFP fluorescence images of 50% (w/v) fluorescein, at different locations of the microfluidic chip to average variation.
2. Load as a stack in Fiji. Take the median through this stack to remove variations between locations (convert to 32-bit output).
   >Image>Stack>ZProject
3. Take a spatial median of the result to get rid of impurities.
   >Process>Filters>Median (Radius= 50 pixels)
4. Subtract a constant, camera-added offset of 100 intensity levels, from the resulting image (100 intensity levels are an arbitrary offset added by the camera to its digitized signal to account for slight negative values).
   >Process>Math>Subtract
5. Divide this image by its median.
   >Process>Math>Divide

The resulting image is the shading corrector.

Use of the shading corrector:

1. Load your image or stack.
2. Subtract the 100 level offset from it.
   >Process>ImageCalculator (Operation: Subtract)
3. Divide this image by the shading corrector (use 32-bit output).
   >Process>ImageCalculator (Operation: Divide)
4. For the resulting image, check the curvature of the fluorescence signal along a line profile from corner to corner – it should be flat.
   >Analyze>PlotProfile

**B. Background subtraction**

Preparation of ‘Mask’ image or stack:

1. Load your shading-corrected GFP image stack, and adjust threshold.
   >Image>Adjust>Threshold
   Adjust the threshold range, choose ‘Default’ and ‘B&W’,
   tick ‘Dark background’, tick ‘Stack histogram’.
   >Apply
   Choose Method: ‘Default’, tick ‘Black background’.

2. Divide the resulting stack by 255.

The pixel values range from 0 to 255. Dividing all the values by 255 will convert it to range from 0 to 1.

>Process>Math>Divide

The resulting stack is the 'Mask' stack.

Use the 'Mask':

1. Load your shading-corrected GFP image stack.
2. Multiply this stack by the 'Mask' stack.

>Process>ImageCalculator (Operation: Multiply)

The resulting stack is ready for image analysis.

**C. Define ROI**

1. Draw ROI around each droplet in one frame.

   >Analyze>Tools>ROI Manager

   Click 'Add' to add all droplet-ROIs into ROI Manager.

   Add also an ROI outside the droplets to confirm the background fluorescence equals to 0.

2. Retrieve data from ROI Manager.

   >More>MultiMeasure

   Tick 'Measure all slices', tick 'One row per slice'.

3. Save the data for further data analysis.

**D. Data analysis**

We analyzed 'Mean' datasets from fluorescence intensity statistics, corresponding to mean gray values, which are the average gray values for each ROI selection. By definition, the mean gray value is the sum of gray values of all the pixels in the selection divided by the number of pixels.

1. Get 'Mean' data for the non-droplet ROI in each position. If it is >0, use these values to additionally correct for background, by subtracting 'Mean' non-droplet fluorescence values from 'Mean' values for the droplet fluorescence signal in each time point.
2. Visually inspect all acquired frames, and exclude the frames in which the entire droplet (or the entire device) was out of focus.
3. Correct time signature by taking into account time from adding phage to the bacterial culture, to the start of the time-lapse experiment (typically, 11-14 min).
4. Correct time signature if there was an acquisition error during the time-lapse experiment, and the acquisition needed to be restarted. Take into account time from the end of the first part of the time-lapse experiment, to the start of the second part of the time-lapse experiment (overall corrections: 31 min for the experiment in **Figure 2A**, 20 min for the experiment in **Figure 3A**).
5. Use the resulting droplet fluorescence intensity values for plots and analysis.
